# Molecular insights into ATP synthase c-ring accumulation in BMP-deficient lysosomes in Batten disease

**DOI:** 10.64898/2026.08.05.743007

**Authors:** Leonhard J. Starke, Erik K. Hansen, Kasparas Petkevicius, Anna L. Duncan

## Abstract

Bis(monoacylglycero)phosphate (BMP) is a signature lysosomal phospholipid that supports the catabolic functions of the lysosome. We recently demonstrated that BMP deficiency is associated with a variant of Batten disease, a neurodegenerative lysosomal storage disorder. This observation led us to investigate how BMP deficiency contributes to the previously reported accumulation of the ATP synthase c-ring in the lysosomes of Batten disease patients and preclinical models. The c-ring is an inner mitochondrial membrane (IMM) protein complex that interacts with cardiolipin, a mitochondrial phospholipid that shares structural features with BMP. Based on this, we hypothesised that BMP may perform an analogous function to cardiolipin in lysosomes. Specifically, we proposed that BMP preferentially interacts with the c-ring, dispersing it within lysosomal membranes and facilitating its degradation. To test this hypothesis, we conducted all-atom molecular dynamics simulations to examine the interactions of various BMP variants with the c-ring of human ATP synthase under different membrane conditions. We observed leaflet-specific preferential interactions of BMP with the protein interface. Replacement of BMP with anionic POPG lipids resulted in a lower binding affinity to the c-ring, indicating the importance of BMP’s unique structure with respect to protein binding. Furthermore, BMP enrichment was enhanced when using the physiologically relevant di-22:6 BMP variant in membranes containing polyunsaturated lipids and cholesterol. Overall, our study suggests that BMP promotes lysosomal c-ring degradation via c-ring co-localisation, whereas BMP deficiency in Batten disease drives c-ring accumulation.

**Significance statement:** Batten disease comprises a group of neurodegenerative lysosomal storage disorders that primarily affects infants and children. Several recently identified disease genes are linked to the biosynthesis of lipid bis(monoacylglycero)phosphate (BMP), a lysosomal lipid required for normal lysosome function. However, how BMP deficiency contributes to disease pathology remains poorly understood. Here, we investigate the relationship between BMP deficiency and lysosomal accumulation of the ATP synthase c-ring, a hallmark pathological feature of Batten disease. Using molecular simulations, we show that BMP preferentially interacts with the c-ring, suggesting that BMP deficiency promotes c-ring aggregation within lysosomes. Our study provides a mechanistic starting point for understanding how BMP deficiency disrupts lysosomal function in Batten disease

## Introduction

Bis(monoacylglycerol)phosphate (BMP) is an anionic phospholipid that is highly enriched in late endosomes and lysosomes.^1,2^ Furthermore, BMP is a structural isomer of phosphatidyl-glycerol (PG), but is distinguished by its unusual stereochemistry and acyl chain arrangement, with both acyl chains attached to glycerol backbones in the S,S configuration, typically at the *sn-2/sn-2’* (2,2’) positions. ^3^ The most widely studied biologically active form contains oleic acid (18:1) bound at the 2,2’ positions (Fig. 1A), although BMP acyl chain composition is tissue-specific and di-docosahexaenoyl (22:6) BMP species are highly enriched in the brain.^4^ The phosphate group of BMP is presumed to be deprotonated under the acidic conditions (pH 4.5–5) inside lysosomes. ^5,6^ This negative charge is linked to a crucial lysosomal function of BMP through the attraction of cationic proteins such as hydrolases. ^3^ BMP is also highly enriched within intralysosomal vesicles, ^2^ and in vitro experiments point towards an essential role for BMP in their formation. ^5^ However, there is still a considerable lack of understanding regarding the detailed functional roles that BMP plays within lysosomes. Importantly, dysregulation of BMP is associated with multiple neurodegenerative diseases, including Niemann–Pick disease^7,8^ and neuronal ceroid lipofuscinosis (NCL).^3^

NCL, also known as Batten disease, refers to a group of severe lysosomal neurodegenerative storage diseases caused by mutations in 14 distinct genes (denoted CLN1–14). ^9^ Batten disease mainly affects children and young adults and is characterised by symptoms such as vision loss and seizures, typically progressing to premature death early in life.^10^ A common hallmark of the disease is the accumulation of a fluorescent pigment called ceroid lipofuscin in various tissue types, including neuronal tissue.^11^ Analysis of the contents of these storage bodies revealed the *c*_8_-ring subunit of mitochondrial ATP synthase as the main protein constituent.^12,13^ While lysosomal ATP synthase subunit c accumulation is widely used as a marker of Batten disease pathology, the molecular mechanisms underlying c-ring accumulation remain poorly understood.

Recently, several CLN genes have been found to be associated with the regulation of BMP levels in lysosomes. ^3^ Specifically, CLN8^4,14,15^ and CLN5^16^ have been identified as essential components of the BMP synthesis pathway. Consistently, rescue experiments on CLN8-deficient animal models show that administration of BMP synthesis precursor restores BMP levels in mice and, crucially, improves neurological phenotypes in zebrafish larvae. ^4^ These results highlight the important causal role of BMP deficiency in Batten disease. ^3,4^

Given the essential role of BMP in Batten disease on the one hand and the accumulation of the *c*_8_-ring subunit on the other, we wanted to investigate whether BMP may be involved in *c*_8_-ring degradation. We were motivated by a potential parallel between BMP and the anionic mitochondrial phospholipid cardiolipin (CL), as both are phosphatidylglycerol (PG)-derived lipids with similar structural features (Fig. 1A). Specifically, solid-state nuclear magnetic resonance (NMR) experiments^17^ and coarse-grained molecular dynamics (MD) simulations ^18^ indicate that CL lubricates the *c*_8_-ring through transient interactions mediated by favourable interactions with a conserved trimethylated lysine residue. These observations led us to hypothesise that BMP may interact with the *c*_8_-ring in lysosomes in a manner analogous to CL, thereby promoting c-ring degradation in the lysosome. To test this hypothesis, we conducted all-atom MD simulations to characterise BMP interactions with the isolated *c*_8_-ring in lysosomal membrane models.

We embedded a human ATP synthase *c*_8_-ring into several membrane models (Table 1) of lysosomal membranes of varying complexity containing either *sn2,sn2’* (S,S’) di18:1 BMP (BMP1), *sn2,sn2’* (S,S’) di22:6 BMP (BMP6), or 16:0–18:1 POPG as a control, and performed unbiased microsecond-timescale atomistic MD simulations. Overall, we find that BMP is enriched at the non-terminal side of the protein and exhibits transient interactions facilitated by the cationic trimethylated lysine residue, similar to the interaction mode previously described for CL.^18^ In contrast, the anionic BMP precursor POPG exhibits a weaker affinity for the protein, as reflected by much shorter interaction times, suggesting that the more exposed phosphate group of BMP allows for tighter binding. Furthermore, simulations of membranes containing cholesterol and polyunsaturated lipids (PUFAs) led to an increase in BMP affinity, particularly for the PUFA BMP variant BMP 22:6-22:6, which is the most abundant BMP species in brain tissue. ^4^

## Methods

### Simulation setup

All-atom MD simulations were performed using the Gromacs software package version 2024^19^] together with the Amber99SB^20^ and Slipids force fields,^21–25^ and the TIP3P water model.^26^ Additional Amber parameters for trimethylated lysine were based on the work of Lu *et al.*,^27^ and parameters for BMP followed those of Enkavi *et al.*^8^

The *c*_8_-ring subunit was retrieved from the PDB database (PDB ID: 8H9F,^28^ chains A–H). Structure preparation was performed using the pdb2gmx command from Gromacs together with custom scripts. The terminal amino acids were treated using the default ionised termini (NH3^+^/COO*^−^*), resulting in a neutral N-terminal aspartic acid and a negatively charged C-terminal methionine. The membrane-core-facing glutamate residue GLU58, which is involved in the proton-transport mechanism of functional ATP synthase (see, for example Blanc *et al.*^29^), was protonated accordingly. Lysine 43 was trimethylated using the corresponding Amber force-field parameters retrieved from http://amber.manchester.ac.uk/ (accessed on 27.04.2026). Both experimental and computational studies indicate that the central cavities of ATP synthase c-rings are filled with lipids.^30–32^ As the cryo-EM structure used here does not resolve lipid densities inside the cavities, we modelled one POPC molecule per leaflet to fill the cavity (Fig. 1G), following our earlier approach^18^ and that of others.^33^ Afterwards, the protein was embedded into each of the five membrane models used throughout this study (Table 1) using the COBY membrane builder.^34^

Lipid topologies and structures for PC, PE, PG, and cholesterol were retrieved from the Slipids lipid database http://www.fos.su.se/∼sasha/SLipids/Downloads.html, and parameters for the *sn2, sn2’* di-18:1 BMP (BMP1) variant were retrieved from https://zenodo.org/records/1034816?. SDPE (18:0/22:6-PE) structure and template charmm topology files were retrieved from CharmmGUI. ^35,36^ The initial partial charges of the SDPE topology files were modified using Amber/Slipids type partial charges for SDPE from existing PE lipids available within the Slipids force field. The structure and topology of *sn2, sn2’* di-22:6 BMP (BMP6) were constructed by modifying the BMP1 tail region using the 22:6 tail from SDPE. Modifications were performed using Pymol^37,38^ to build the initial coordinate files and Gromologist^39^ to generate the Gromacs topology files. The modified lipids structures were energy-minimised and equilibrated for 5 ns to relax the initial structures. To allow dense packing of lipids into membranes using COBY, all lipid input structures were preprocessed using the MemGen webserver,^40^ resulting in more compact lipid structures.

**Table 1:** Membrane compositions of the 5 studied systems.

| System | POPC | POPE | DOPC | DOPE | SDPC<br>(18:0/22:6) | SDPE<br>(18:0/22:6) | BMP1<br>(di 18:1) | BMP6<br>(di 22:6) | POPG | CHL1 |
| --- | --- | --- | --- | --- | --- | --- | --- | --- | --- | --- |
| noBMP | 66 % | 33 % |  |  |  |  |  |  |  |  |
| baseBMP1 | 55 % | 25 % |  |  |  |  | 20 % |  |  |  |
| basePOPG | 55 % | 25 % |  |  |  |  |  |  | 20 % |  |
| BrainBMP6 |  |  | 15 % | 15 % | 15 % | 15 % |  | 20 % |  | 20 % |
| BrainBMP1 |  |  | 15 % | 15 % | 15 % | 15 % | 20 % |  |  | 20 % |

The membrane builder COBY was initially developed for coarse-grained systems, and although the tool is in principle agnostic with respect to the type of input structures, additional steps were required to ensure stable atomistic simulations of the *c*_8_-ring in heterogeneous membranes. Therefore, the following extended equilibration protocol was applied: (1) Before solvation, the generated protein–membrane system was energy-minimised in vacuum using a soft-core potential to remove potential steric clashes between atoms. This was followed by a second short energy minimisation with the soft-core potential switched off. As each system contains a net charge, neutralising ions were added to avoid introducing additional artefacts during minimisation.^41^ During minimisation and subsequent equilibration, position restraints were applied to the protein backbone and heavy side-chain atoms, as well as to the phosphorus and hydroxyl oxygen atoms of the phospholipid and cholesterol headgroups. Additional dihedral restraints were applied to the glycerol linkers and double bonds in the lipid tails. (2) The minimised systems were solvated with TIP3P water, and Na^+^ and Cl*^−^* ions were added to reach an ion concentration of 150 mM. (3) The solvated systems were subsequently energy-minimised and equilibrated following a protocol consisting of six short equilibration simulations with increasing time steps and progressively reduced force constants on the position restraints to relax the systems. A summary of the main settings used during the equilibration protocol is provided in Table S1.

After equilibration, production runs were set up using hydrogen mass repartitioning^42^ with a mass-repartitioning factor of 2.5, as recommended by Jung *et al.*^43^ for AMBER force fields. Systems containing cholesterol were simulated with a time step of 3 fs to ensure numerical stability, whereas all other systems were simulated using a 4 fs time step. The temperature was set to 300 K and controlled using the velocity-rescaling thermostat.^44^ The pressure was set to 1 bar using semi-isotropic pressure coupling with the c-rescale barostat^45^ during both equilibration and production runs. Water hydrogen bonds were constrained using SETTLE,^46^ whereas all other hydrogen bonds were constrained using LINCS.^47^ Van der Waals interactions were described using a Lennard–Jones potential with a cut-off of 0.9 nm. Coulomb interactions were calculated using the PME approach.^48^ For each membrane system, five independent replicas were generated and simulated for 12 µs, except for the noBMP system, where we simulated 3 replicas of 5 µs and 2 replicas for 6 µs.

### Basic analysis of MD simulations

MD simulation analysis was performed using custom scripts together with the software packages Gromacs,^19^ MDAnalysis,^49,50^ PyLipID,^51^ and mosaics.^52^ Structures were visualised using VMD 2.0^53^ https://www.ks.uiuc.edu/Research/vmd/ and Pymol^37,38^ .

Trajectories were preprocessed by centring the protein within the simulation box and removing protein rotation in the xy-plane. The first 500 ns of the production runs were discarded to ensure that no potential artefacts from the system-building procedure remained. The POPC lipids within the central cavity were excluded from the analysis. Unless otherwise stated, analyses were conducted separately for the upper and lower membrane leaflets. Leaflet assignment of the phospholipids was performed using the MDAnalysis LeafletFinder algorithm. As cholesterol exhibited frequent leaflet flip-flop, its analysis was performed for the membrane as a whole.

**Density calculation:** The volmap tool in VMD was used to visualise lipid densities around the protein for one representative trajectory of the baseBMP and noBMP membrane models. Hereby, densities were calculated based on the positions of the phosphorus headgroup atoms using a grid spacing of 1 Å. Densities were visualised using an isovalue cut-off of 0.0014 1/ Å^3^.

**Root-mean-square deviation (RMSD):** RMSD traces were calculated for all trajectories using the protein backbone atoms and the Gromacs tool rmsdist. Hereby, the first frame of each trajectory was used as the reference structure.

**Depletion–enrichment (DE) index:** The DE index for a given lipid species *l_i_* is defined via^54^

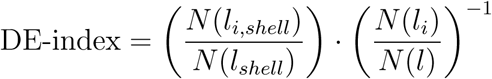

Whereby, *N* (*l_i,shell_*) and *N* (*l_shell_*) are the numbers of lipids from species *l_i_* and the total number of lipids within a cut-off-defined protein shell, respectively. *N* (*l_i_*) is then the total number of species *l_i_* and *N* (*l*) the total number of lipids. Here, we used a cut-off distance of 0.7 nm based on the distance between the phosphorus atom of the lipid headgroup and the nearest protein atom. DE indices for cholesterol were computed for both membrane leaflets together due to frequent flip-flop events, using the oxygen atom of the hydroxyl headgroup as the reference.

**Membrane thickness:** To evaluate the position-dependent membrane thickness, 2D membrane-thickness maps were calculated using the mosaics software suite.^52^ Hereby, membrane thickness was inferred from the time-averaged z-positions of the phospholipids using the glycerol backbone atoms as reference. Subtracting the averaged z-positions from each other yielded one 2D thickness map per trajectory. For each map, a 1D thickness profile as a function of the distance from the protein centre was obtained by radial averaging.

### Analysis of lipid residence times and occupancies

Interfacial lipid–protein interactions were analysed using the PyLipID package. ^51^ Hereby, lipid–protein contacts were determined using a dual cut-off scheme, in which a contact was considered established when the lipid moved within the inner cut-off distance and terminated once the lipid moved beyond the outer cut-off distance. Here, we used 0.5 and 0.8 nm for the inner and outer cut-offs, respectively. To reduce computational cost and allow potential comparison with coarse-grained simulations, pseudo-coarse-grained trajectories were generated containing only the phosphorus atoms of the phospholipid headgroups and pseudo-coarse-grained beads for the amino acids, defined by the centre of geometry of groups of heavy atoms. The mapping of heavy atoms followed the scheme established for Martini 3 proteins.^55^ Lipid–protein distances were evaluated every 20 ns for simulations using a 4 fs time step and every 30 ns for simulations using a 3 fs time step for each phospholipid species and membrane leaflet. This coarse temporal sampling was chosen to focus on long-timescale lipid-binding events and to facilitate comparison with coarse-grained simulations. Lipid–protein contacts were subsequently defined using the described cut-off scheme.

By default, PyLipID clusters nearby contacting protein residues to define local lipid-binding sites along the protein interface. In contrast, we defined a single binding site per membrane leaflet. This definition allows the inclusion of lipid-binding events in which lipids move along the protein surface. In addition, lipid contacts were classified by protein chain, and time traces were generated for each contacting lipid with examples for BMP1 and POPE are shown in Figure S3,S4. Lipid contact durations were pooled and sorted to calculate the contact survival function *σ*(*t*), as detailed in.^51^ An average residence time was then obtained by ordinary least-squares fitting of the survival function *σ*(*t*) to a biexponential model function defined as

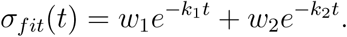

Here *k*_1_, and *k*_2_ are the rates describing the decay of lipid contacts, while *w*_1_ and *w*_2_ are the weights describing the population of the corresponding decay processes (*w*_1_ + *w*_2_). By default, PyLipID assumes that *σ_fit_*(*t*) exhibits a distinct slow and a fast decay mode and defines the residence time via *τ* = 1*/*(min(*k*_1_*, k*_2_)). This implicitly assumes that the smaller *k* corresponds to the larger weight. However, we found that this condition was not fulfilled for some simulation systems. Therefore, we instead calculated a mean residence time defined via

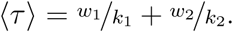

The uncertainty of the mean residence time ⟨*τ* ⟩ was calculated using the uncertainties of the fitted parameters obtained from the covariance matrix of the fit. This procedure was found to be more robust than the original implementation, which uses bootstrapping of the survival function to estimate fit uncertainties. We note that, ideally, one should relax the assumption of two decay modes and instead use a Bayesian approach to infer the residence-time scales, as described by Sexton et al. ^56^ However, this would require substantially more exhaustive sampling on the order of at least approximately 100 µs, which would likely necessitate the use of coarse-grained force fields and thus sacrifice atomistic resolution.

Lipid occupancies were calculated per residue as the fraction of frames in which the residue was in contact with a lipid of the specified lipid type. Occupancies were then averaged over the eight identical chains to calculate occupancy-per-residue histograms. To measure the percentage of the contact time for 1 vs 2 protein chains we calculated for each lipid contact *c_i_*, the time Δ*t*(*c_i_, n_c_*) during which the contact was formed between a lipid and a specific number of chains (*n_c_* = 1, 2). The contact time percentage *t_c_* for a given *n_c_* is then defined as

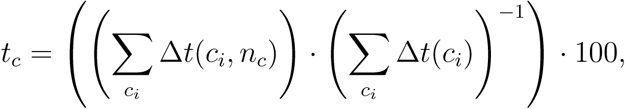

with Δ*t*(*c_i_*) denoting the overall duration of the contact.

**Figure 1:**
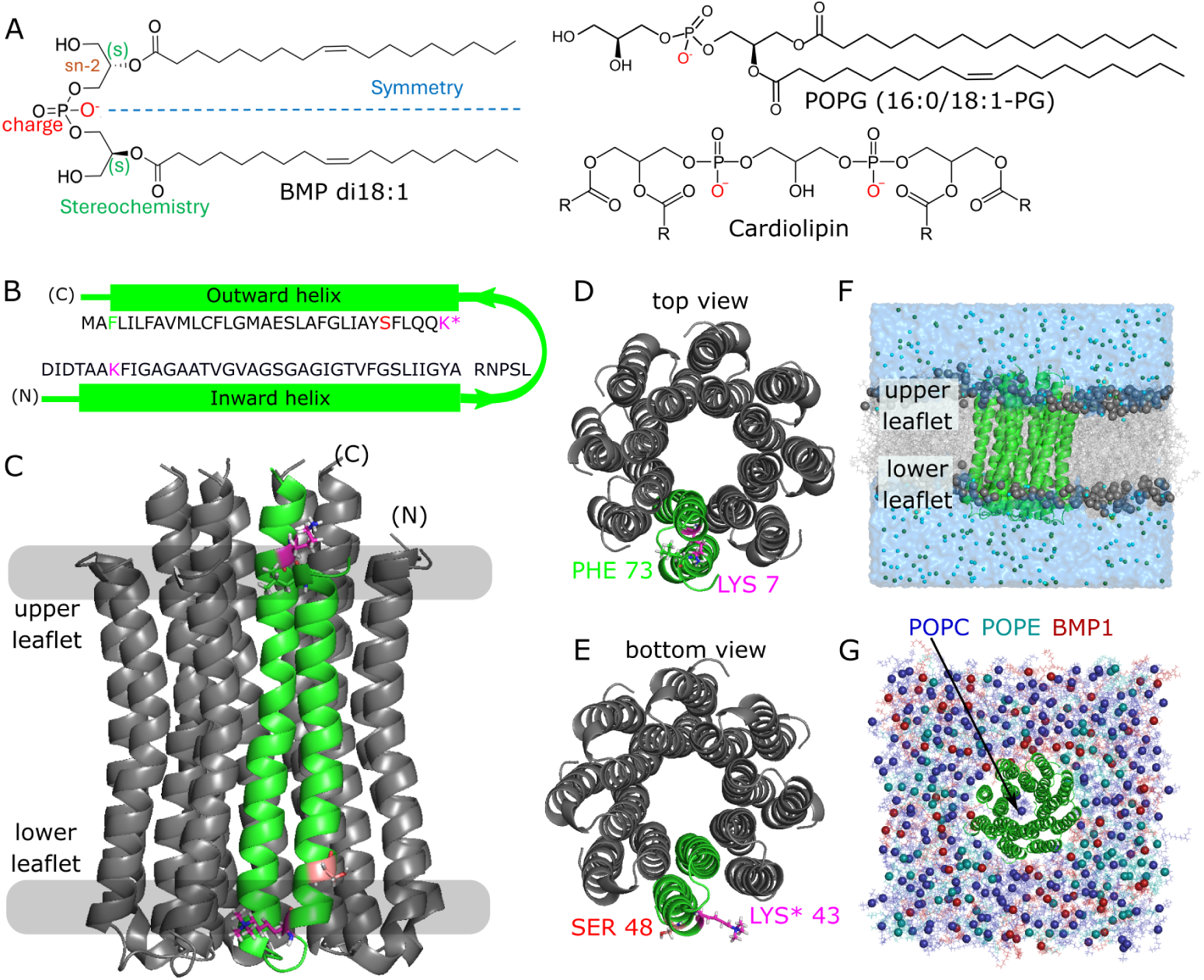
Structure of BMP1 and the ATP synthase *c*_8_-ring. (A) Left panel: Chemical structure of di-18:1, *2S,2S’* BMP. The symmetry axis, charge, and stereochemistry are highlighted. Right panel: Chemical structures of 16:0-18:1 PG (POPG) and generic cardiolipin (R indicates position of fatty acid hydrocarbon chains). The locations of negative charges are indicated in red. (B) Amino acid sequence and secondary structure of the human ATP synthase *c*-subunit. Selected residues LYS7, trimethylated LYS43 (LYS*43), SER48, and PHE73 are highlighted in colour. (C–E) Cartoon representation of the human *c*_8_-ring (PDB ID: 8H9F^28^), with one chain highlighted in green, N and C termini indicated for one chain and selected residues shown in stick representation. (C) Side view indicating the lipid phosphate group positions as grey area with the designated upper and lower leaflets indicated. (D,E) Top and bottom views. (F) Snapshot of the baseBMP simulation system (final simulation frame) shown from the side (Table 1). The protein (green) is shown in cartoon representation, lipids are shown in grey with phosphorus atoms represented as spheres, ions are shown as small spheres (Na^+^, cyan; Cl*^−^* dark green), and water is shown as a transparent blue surface. (G) Top view of the simulation system with solvent removed for clarity. Lipid phosphorus atoms are coloured according to lipid species (dark blue, POPC; teal, POPE; dark red, BMP1). The arrow indicates one POPC molecule per leaflet in the central cavity.

## Results

### Structure of the human *c*_8_-ring in the lysosomal membrane

In this study, we examined BMP interactions with a human mithochondrial ATP synthase *c*_8_-ring based on a cryo-EM structure reported by^11^Lai et al.^28^ (PDB ID: 8H9F). In mitochondria, the c-subunit of the membrane-embedded F_0_ domain of ATP synthase functions as the main rotational unit driving ATP synthesis.^29,57^ Each individual c-subunit consists of two *α*-helices connected by a short loop, forming a hairpin transmembrane domain (Fig. 11). In humans, eight *c*-subunits form an *α*-helical barrel, with the longer N-terminal helix forming the inner ring around the small central cavity and the shorter C-terminal helix forming the outer ring, which constitutes the main interface with the surrounding membrane (Fig. 1C–E).

A conserved membrane-facing trimethylated lysine^58^ (LYS*43), located on the non-terminal side of the protein close to the loop region (Fig. 1E), has been suggested to be involved in the accumulation of cardiolipin at the *c*-ring, as observed in our previous coarse-grained MD simulations. ^18^ This is consistent with experimental studies of bacterial *c*-rings exhibiting clear preferences for cardiolipin.^17,59^ In addition, we previously identified preferential interactions of cardiolipin with a nearby serine residue (SER48) (Fig. 1C, E), as well as with a lysine (LYS7) and phenylalanine (PHE73) on the terminal side of the protein (Fig. 1C, D). We modelled one POPC per leaflet into the the central cavity of the *c*_8_-ring (Fig. 1G), as previous experimental and computational studies indicated that the cavity is filled with lipids^30–32^ (see Methods).

The *c*_8_-ring was embedded in a membrane composed of 55% POPC, 25% POPE and 20% BMP1, hereafter referred to as the baseBMP membrane model (Fig. 1G, Table 1). The protein structure does not exhibit significant conformational changes throughout the simulations, as exemplified by overall backbone root-mean-square deviations within 0.2 nm (Fig. S1).

### BMP1 is selectively enriched at the *c*_8_-ring

To assess whether BMP exhibits preferential interactions with the *c*_8_-ring, we performed MD simulations of the *c*_8_-ring embedded in the baseBMP membrane model and compared them to simulations using a membrane model containing 67% POPC and 33% POPE, hereafter referred to as the noBMP membrane model (Table 1). Examination of the lipid headgroup density using the volmap tool in VMD showed that BMP1 is enriched at the protein interface within the lower membrane leaflet, whereas no BMP1 enrichment was observed in the upper leaflet (Fig. 2A, left panel). For the system without BMP1, no strong enrichment of either POPC or POPE was observed (Fig. 2A, right panel). To quantify lipid enrichment, we calculated the depletion–enrichment index (DE-index)^54^ for each lipid species in both leaflets of both simulation systems, using the minimum distance between the lipid headgroup phosphorus atom and the protein to define lipid–protein contact. Examination of the DE-index distributions confirmed the observations from the volmap analysis, with BMP1 being the only lipid significantly enriched at the protein interface, and this enrichment being limited to the lower membrane leaflet (Fig. 2B).

**Figure 2:**
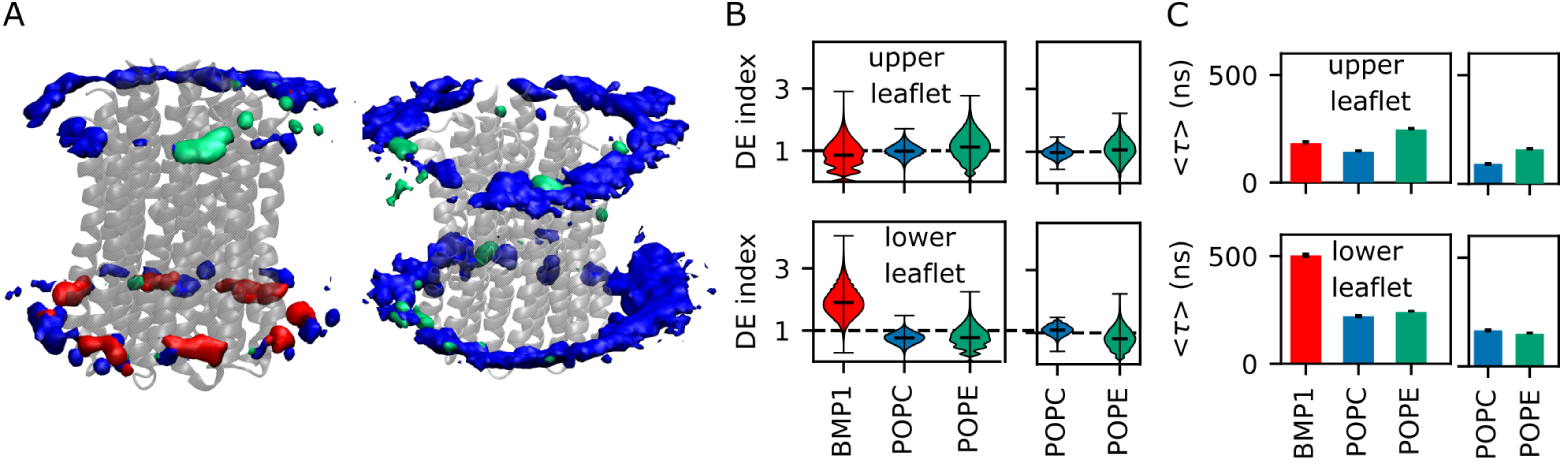
Lipid–protein interactions in simple membrane systems with and without BMP1. (A) 3D density of the phosphorus atoms of BMP1 (red), POPC (blue), and POPE (turquoise) around the *c*_8_-ring (transparent grey, shown in cartoon representation) shown for representative trajectories of the baseBMP system (left) and the noBMP system (right). Densities were obtained using the volmap tool in VMD (isovalue 0.0014 1/Å^3^, grid size 1 Å). (B) Depletion-Enrichment (DE)-index distributions for each leaflet and lipid species. Distributions, as well as median, maximum, and minimum values, are shown in black. The dotted line shows DE= 1, which indicates no lipid enrichment at the protein, relative to bulk lipid composition. (C) Mean residence times for each leaflet and lipid species. Error bars were calculated by error propagation of the fit-parameter uncertainties.

To estimate the affinity of each lipid for the protein, we calculated lipid contact durations using the distance between the lipid headgroup phosphorus atom and the protein to construct a contact survival function *σ*(*t*), using PyLipID^51^ (Fig. S2). By fitting the survival functions to a biexponential model, we obtained a mean residence time ⟨*τ* ⟩ for each lipid species in each leaflet and membrane system. The residence times are consistent with the information obtained from the DE-index, with BMP1 in the lower membrane leaflet exhibiting longer residence times compared to all lipids in the upper membrane leaflet (Fig. 2C). In the POPC/POPE membrane system, both lipids exhibit comparatively short residence times, similar to their behaviour in the BMP1-containing membrane system (Fig. 2C). Thus, BMP1 exhibits an increased leaflet-specific affinity, leading to a relatively homogeneous accumulation around the *c*_8_-ring (Fig. 2A). Overall, lipid contacts appear to be transient, with only a few long-lasting BMP1 contacts observed in the lower membrane leaflet (Fig. S3 C).

**Figure 3:**
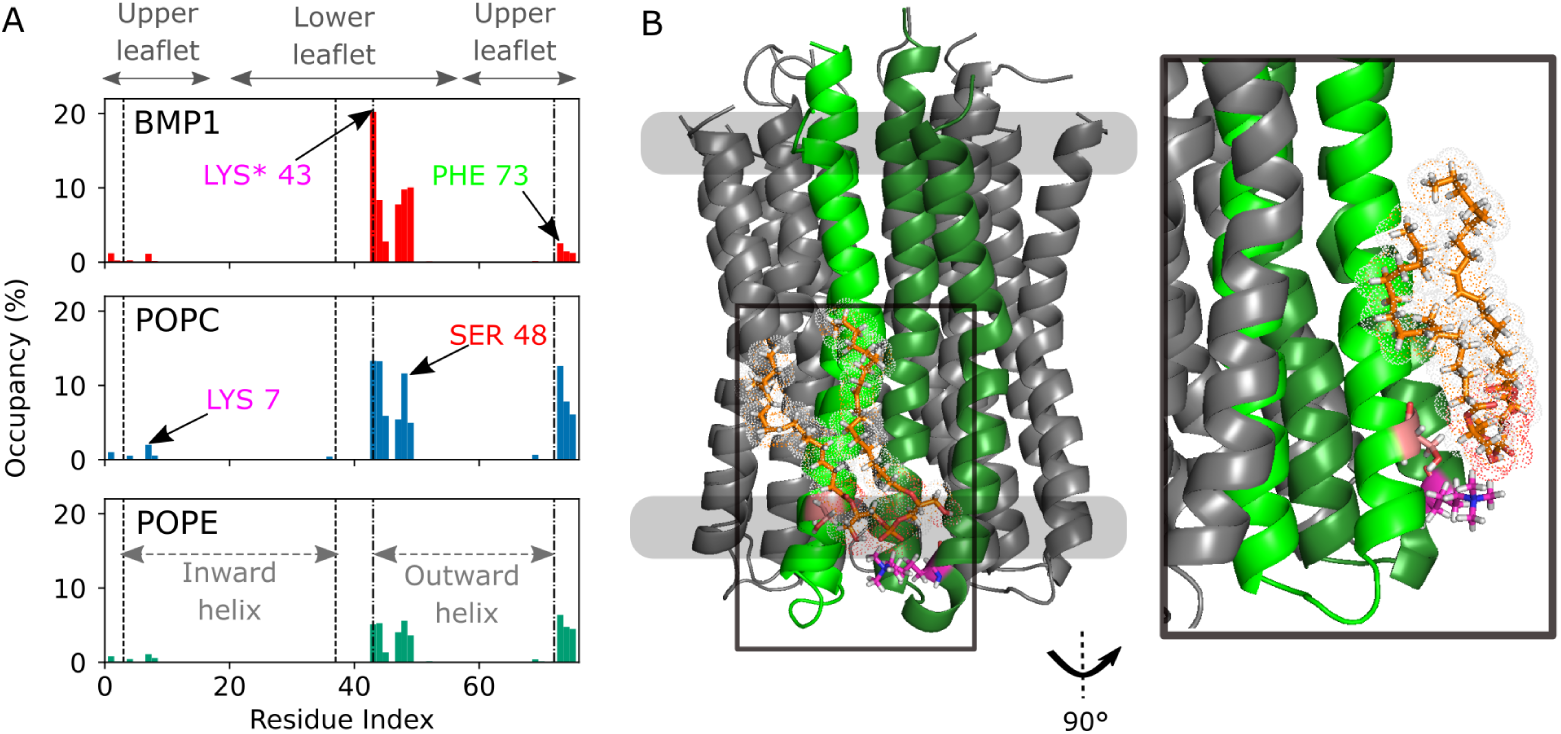
Preferential lipid–protein residue interactions. (A) Averaged lipid occupancies per protein residue for BMP1, POPC, and POPE. The approximate positions of the membrane leaflets and the start and end points of the helices are indicated as a guide to the eye. Specific residues corresponding to the highest occupancies are highlighted. (B) Example snapshot of a bound BMP1 molecule at the protein interface in the lower membrane leaflet (stick representation with carbon atoms coloured orange). Contacted chains are indicated in green and dark green, and the interacting residues LYS*43 and SER38 are highlighted using the same representation as in Fig. 1. The inset shows a close-up side view of the lipid in stick-and-dot representation.

### BMP1 preference is mediated by headgroup charge-charge interactions

To examine which residues are involved in lipid–protein contacts, we used PyLipID to calculate occupancies for each lipid species on a per-residue basis. Figure 3A shows the lipid occupancy per residue averaged over the eight protein chains. The highest lipid occupancies are found on the non-terminal side of the protein around residues 43–45 and 47–49, as well as at the C-terminal residues 72–75. In contrast, the N-terminal residues, including LYS7, exhibit only low lipid occupancies (Fig. 3A). This likely reflects their positioning behind the outer helix and consequently their lower exposure to lipid headgroups. More importantly, BMP1 occupancy in the lower membrane leaflet exhibits a pronounced peak at LYS*43, whereas POPC and POPE exhibit similar occupancies for LYS*43/GLN44 and SER48. This suggests that BMP1 headgroup forms preferential charge–charge interactions, which likely lead to the observed enhanced affinity compared to POPC and POPE.

Visual inspection of bound BMP1 reveals a binding pose in which the BMP1 headgroup is positioned further towards the hydrophobic core of the membrane, relative to LYS*43 side chain, allowing the lipid to occupy the small groove between two neighbouring chains (Fig. 3B). To quantify the importance of this binding mode and compare it with POPC and POPE, we determined the contacted protein chains for each lipid contact at every simulation time point. An example time trace for BMP1 and POPE from a single trajectory is shown in Fig. S3 and S4 respectively. Examination of the time traces showed that, during a single contact event, lipids interact mostly with one or two neighbouring chains. However, there are also cases where a lipid translocates from one protein groove to the next, as indicated by lipids contacting three to four neighbouring chains within a single trajectory (cf. lipid no. 5 in Fig. S3 C, lipid 1 in Fig. S4 B). To quantify the frequency of lipid contacts involving multiple chains, we calculated the percentage of time a lipid stays in contact with 1 or 2 chains for each lipid species and leaflet (Fig. S5). We found that BMP1 headgroups in the lower membrane leaflet are more frequently in contact with two adjacent subunits compared to POPC and POPE, potentially allowing BMP1 to form tighter interactions with the protein interface.

### POPG exhibits reduced binding affinity to the *c*_8_-ring

Given that the preferential charge–charge interactions of BMP1 with LYS*43 are likely responsible for the lipid accumulation at the *c*_8_-ring, we wanted to examine whether this is a general feature of *c*_8_-ring interactions with anionic lipids. In order to explore this further, we ran further simulations replacing BMP1 with POPG, a closely related anionic phospholipid and biosynthetic precursor of BMP. Thus, we performed simulations of the *c*_8_-ring embedded in a membrane containing 55% POPC, 25% POPE, and 20% POPG, hereafter referred to as the basePOPG membrane model, and calculated DE-indices, mean residence times ⟨*τ* ⟩, and residue occupancies as shown in Figure 4. We again observed enrichment of POPG in the lower membrane leaflet (Fig. 4A); however, the residence time distribution for POPG is characterized by a higher probability of shorter residence times compared to BMP1 (Fig. S6 A) resulting in a overall lower ⟨*τ* ⟩ (Fig 2C,Fig. 4B). Consistently, POPG exhibits an occupancy distribution more similar to that of POPC and POPE with a preferential interaction at SER48 (Fig. 4D). The lower affinity of POPG highlights the importance of the specific stereostructure of BMP1 and its exposed phosphate headgroup. In contrast, POPG’s bulky glycerol headgroup partly shields the negative charge, as exemplified by a representative snapshot of POPG bound to the *c*_8_-ring (Fig. 4C). Consistently, the contacted time for POPG is shifted towards a larger fraction of single-chain contacts (Fig. S5 B).

### BMP *c*_8_-ring interactions are enhanced by increased membrane thickness and lipid unsaturation

Finally, we wanted to examine a physiologically relevant scenario and therefore conducted simulations using more complex lipid compositions, hereafter referred to as the brainBMP1 and brainBMP6 membrane models (Table 1). Hereby, we accounted for the presence of cholesterol at intermediate concentrations, ^60^ as well as the abundance of docosahexaenoic acid (DHA, 22:6) acyl chains in brain tissue. ^61^ Specifically, we selected 18:0/22:6 PC and PE (SDPC/SDPE), as this acyl-chain combination is enriched in neurons and considered to be functionally relevant.^62^ Furthermore, we performed simulations using both the brainBMP1 membrane model containing BMP1 and the brainBMP6 membrane model containing *sn2,sn2’* di-22:6 (S,S’) BMP (BMP6), with the latter representing the dominant brain BMP species. ^4^ The DE-indices and residence times of the phospholipids in both membrane compositions (Fig. 5A-B) exhibit the same trends as observed for the simpler membrane systems. Both BMP species are enriched only in the lower membrane leaflet (Fig. 5A), exhibit pronounced interactions with LYS*43 (Fig. 5C), and display notably longer residence times compared to the PC and PE species (Fig. 5B). Furthermore, BMP6 exhibits slightly longer residence times (Fig. 5B, Fig. S6 B) compared to BMP1, which is reflected by BMP6’s higher occupancy maximum at LYS*43 (Fig. 5C). Thus, the results are overall consistent, with the marginally higher BMP6 affinity indicating a possible functional role of the unusually high unsaturation of its acyl tails.

**Figure 4:**
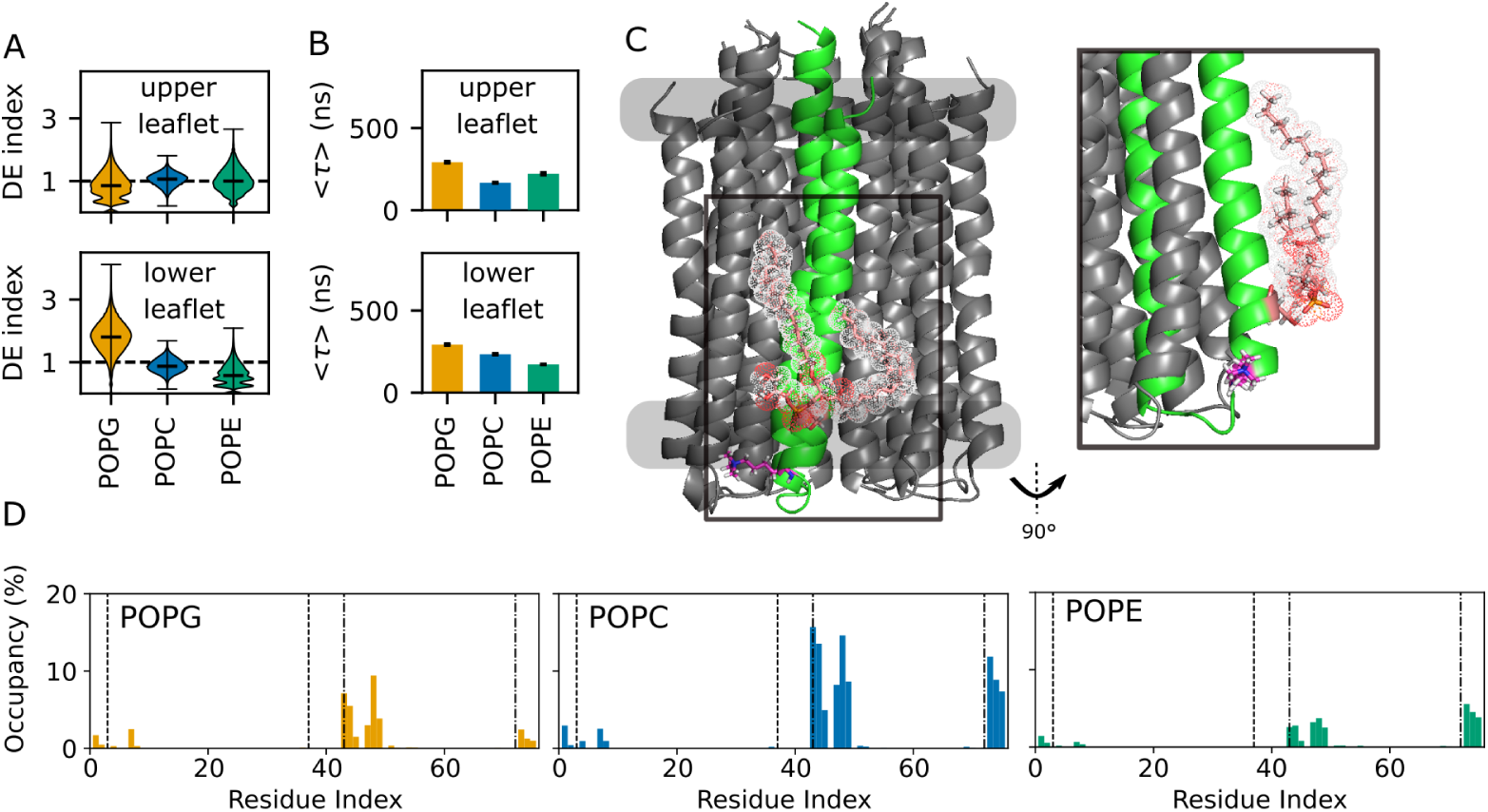
Interactions of POPG-containing membranes with the *c*_8_-ring. (A) DE-indices and (B) average residence times shown for each lipid species and membrane leaflet (C) Snapshot of POPG (stick-and-dot representation with carbon atoms coloured light red) with the headgroup interacting primarily with a single *c*_8_-ring subunit, again highlighting LYS*43 (magenta) and SER48 (light red). The inset shows a zoomed-in view of the POPG headgroup interacting with SER48. (D) Averaged lipid occupancies per protein residue for each lipid species.

One potential explanation for the observed small difference in BMP affinities may be provided by distinct interactions with cholesterol in the studied membrane systems. As trajectory inspection revealed frequent cholesterol flip-flop events between the membrane leaflets, we calculated the DE-index of cholesterol for the membrane as a whole. We found that cholesterol is not enriched in either system and, in the presence of BMP6, appears to be slightly depleted from the protein interface (Fig. S7 A). As cholesterol is known to be depleted from highly unsaturated membrane domains,^63^ it is possible that BMP6 accumulation at the *c*_8_-ring and cholesterol depletion are coupled, leading to the observed increase in BMP6 affinity for the protein.

Another important aspect to consider is the role that membrane composition plays in modulating membrane structure, specifically with respect to cholesterol, which is known to increase membrane order and consequently membrane thickness.^64,65^ We therefore calculated 2D membrane-thickness maps from all simulations and plotted the radially averaged thickness profiles to determine the membrane thickness as a function of distance from the protein (Fig. S7 B). Overall, the membrane thickness in the membrane bulk phase is substantially larger for cholesterol-rich membranes, consistent with previous reports.^64,65^ More importantly, all membrane systems exhibit a visible thickening at the protein rim, indicative of a strong hydrophobic mismatch between the membrane and the protein. While the relative increase in thickness is strongest for the systems without cholesterol, the greatest membrane thickness at the protein rim is observed for the cholesterol-containing membrane systems. The observed overall increase in membrane thickness for the brain-mimicking membranes may contribute to the enhanced affinity of BMP for the *c*_8_-ring by enhancing the presentation of the BMP headgroup at the membrane interface and thereby reducing the distance to the charged LYS*43.

**Figure 5:**
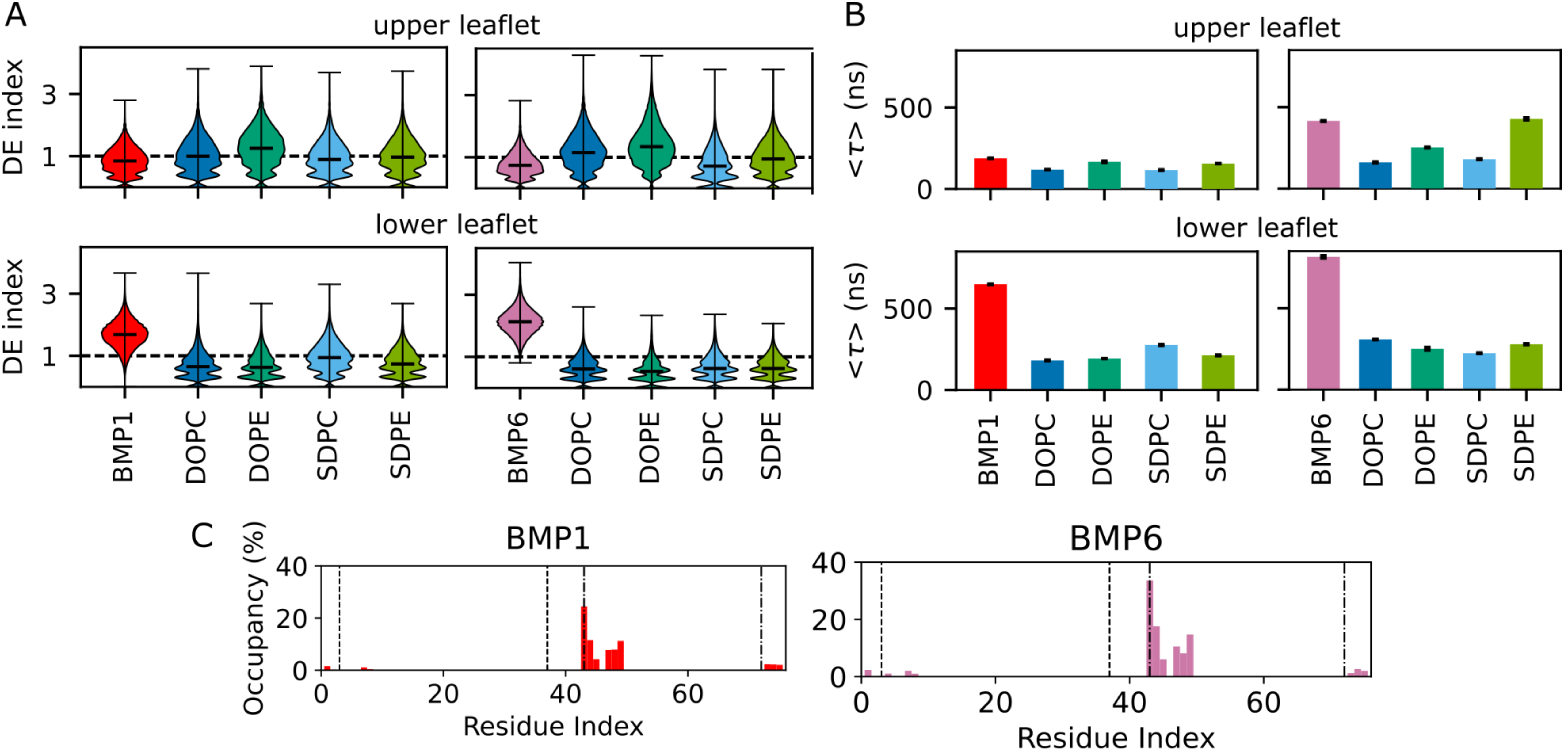
BMP *c*_8_-ring interactions in complex membranes. (A) Comparison of phospholipid DE-index distributions for the brainBMP1 membrane model (left) and the brainBMP6 membrane model (right). (B) Comparison of average residence times for each membrane leaflet and phospholipid species in the brainBMP1 membrane model (left) and the brainBMP6 membrane model (right). (C) Comparison of average occupancies per residue for the brainBMP1 membrane model (left) and the brainBMP6 membrane model (right).

## Discussion

In this study, we conducted all-atom MD simulations of lipid interactions with the human ATP synthase *c*_8_-ring, focusing on interactions with the lysosomal anionic phospholipid BMP. We found that anionic BMP is selectively enriched around the positively charged trimethylated LYS*43 residues. This leads to the accumulation of BMP in the lower membrane leaflet around the c-ring and to increased residence times compared to zwitterionic PC/PE lipids and anionic PG. Analysis of lipid binding poses in the LYS*43-proximal membrane leaflet indicates that the increased affinity of BMP for the *c*_8_-ring is caused by its exposed phosphate headgroup, which allows it to position itself above the LYS*43 side chain (Fig. 33B). This is also reflected by the shift in the contacted-chain occupancy of BMP (Fig. S5) towards positions within the grooves formed by two adjacent chains. Furthermore, BMP affinity for the *c*_8_-ring is enhanced in more native-like membranes containing cholesterol and PUFA phospholipids. We suggest that the observed increase in membrane thickness at the membrane–protein interface (Fig. S7 B) may enhance BMP headgroup interactions with LYS*43. Consistently with the high abundance of PUFA acyl chains in the brain, the highest observed affinity was found for the physiologically relevant BMP6 variant (Fig. 5C, Fig. S5 B).

The observed BMP interactions with the human *c*_8_-ring resemble the interaction pattern previously reported for CL interactions with c-rings by Duncan *et al.* using coarse-grained MD simulations. In both cases, the lipids form transient interactions with the protein and exhibit comparable residence times of approximately 600 ns, mediated largely by lysine residues. However, the previous study also found CL enrichment at the terminal side of c-rings facilitated by LYS7, whereas we did not observe a pronounced interaction of BMP with LYS7 (Fig. 3A). These differences may arise due the use of the coarse-grained Martini 2 force field^66^ compared to our atomistic model. At the same time, they likely indicate a qualitative difference in lipid–protein interactions between CL and BMP, despite both lipids being characterised by their exposed phosphate headgroups (Fig. 1A).

Other examples of BMP–protein interactions include an atomistic computational study by Enkavi *et al.*,^8^ which examined BMP1 interactions with the peripheral membrane protein NPC2. In that study, tight NPC2 membrane association required BMP specifically, as substitution with DOPG resulted in weaker binding compared to BMP. In addition, the authors observed that cholesterol facilitates the “presentation” of BMP at the membrane interface, thereby enhancing protein binding. Moreover, a combined experimental and computational study of BMP interactions with the 3TAT peptide^67^ observed a similar preference for the BMP headgroup to be exposed at the membrane surface, leading to more pronounced interactions with the peptide compared to POPG. Furthermore, the study found higher binding of 2,2-BMP to the peptide compared to 3,3-BMP. While these results are not directly comparable to our case, given that they describe peripheral membrane-protein binding, they nevertheless highlight the specific structure of BMP as a potentially key component underlying its crucial role within lysosomal and endosomal membrane systems.

We should note that our computational model has several limitations. First, our brain-mimicking membrane compositions (brain BMP1 and brainBMP6) represents only a proxy for the true lysosomal membrane composition in brain in vivo. Second, atomistic simulations of lipid–protein interactions provide only limited sampling of rare long-timescale lipid-binding events. For instance, we observed a single 4 µs binding event of POPC in the POPG-containing membrane system at the lower leaflet (Fig. S2 C), which introduces a strong bias into the residence-time calculation when following the approach outlined in the original PyLipID publication.^51^ Instead, we calculated a mean residence time as a proxy for membrane affinity. Nevertheless, our results demonstrate that atomistic simulations can resolve qualitative differences in lipid–protein interactions for moderately sized membrane proteins. This allowed us to study the enrichment of BMP at the *c*_8_-ring, suggesting a link between *c*_8_-ring accumulation and BMP deficiency in Batten disease.

Previous research on c-ring accumulation in Batten disease has identified disease-gene products involved in protein degradation. Specifically, the CLN2 gene product TPP-1 was described as the primary protease responsible for initiating cleavage of the *c*_8_-ring subunits.^68^ As BMP is thought to recruit proteases to the lysosomal membrane, one can speculate that its enrichment at the *c*_8_-ring allows for targeted cleavage of, for example, the c subunits. Furthermore, BMP may also contribute to stabilising TPP-1 and other proteases against self-degradation within the lysosome. Further studies are therefore needed to investigate the molecular interactions of TPP1 and other proteases, such as cathepsin D, with the *c*_8_-ring in the presence of BMP.

Collectively, our simulations suggest that BMP forms specific interactions with the *c*_8_-ring and thereby promote its degradation within lysosomal membranes. These findings support a model in which BMP facilitates lysosomal *c*_8_-ring degradation through increasing its accessibility to lysosomal proteases, whereas BMP deficiency in Batten disease promotes *c*_8_-ring accumulation.

## Supporting information

The supporting material contains supporting figures S1-S7 and table S1.

## Data availability

The MD simulation setup files and trajectories for all simulation systems as well as the inhouse modified version of pylipID are freely available. Simulation setups and trajectories for the noBMP,baseBMP & basePOPG system and the pylipID code are deposited at: https://zenodo.org/records/20793991. Simulation setups and trajectories for the brainBMP1/BMP6 systems are deposited at: https://zenodo.org/records/20796969. COBY is publicly available under: https://github.com/MikkelDA/COBY.

## Author Contributions

L.J.S designed and performed research, analysed data and wrote the manuscript. E.K.H performed research and reviewed and edited the manuscript. K.P. conceptualized and designed the research and reviewed and edited the manuscript. A.L.D. designed the research, provided software, supervised the project, reviewed and edited the manuscript and acquired funding.

## Acknowledgments

The authors acknowledge funding support from the Danmarks Frie Forskiningsfond via the Inge-Lehman grant (2098-00023B), UKRI Medical Research Council (MC_UU_00028), Academy of Medical Sciences (SBF0010/1078). Computational resources were provided by the Grendel cluster of the Centre for Scientific Computing Aarhus with funding provided by the Resource for Biomolecular Simulations (ROBUST; supported by the Novo Nordisk Foundation; NNF24OC0087976). L. J.S acknowledges Mikkel D. Andreasen and Amanda Stange for their help with the simulation setup workflow, Miłosz Wieczór for support with respect to the lipid topology preparation and Mario Vazdar and Jean-Philippe Pellois for helpful feedback with respect to BMP1 parameters. E.K.H acknowledges support from the Erasmus+ Internship Funding programme through RWTH Aachen University, which enabled the research stay abroad associated with this work. The programme did not provide direct funding for the research activities reported in this publication. We also thank Prof. Sir John E. Walker and Prof. David Palmer for valuable discussions that helped shape the conceptual development of this work.

## Declaration of generative AI and AI-assisted technologies in the writing process

ChatGPT (OpenAI) was used to assist with language refinement, and the authors take full responsibility for the final content.

