## Supplementary material for "Molecular insights into ATP synthase c-ring accumulation in BMP-deficient lysosomes in Batten disease": The supporting material contains supporting figures S1-S7 and table S1.

**Table S1** Overview of minimization and equilibration protocol, detailing number of simulation steps, time step, force constants (Fc) for protein Backbone (bb) and sidechains (sc), lipids phosphorous atom (l), lipid dihedrals (d).

| Step | $N_{steps}$ | Δt (fs) | Fc,bb (kJ/mol nm$^2$) | Fc,sc (kJ/mol nm$^2$) | Fc,l (kJ/mol nm$^2$) | Fc,d (kJ/mol nm$^2$) |
|---|---|---|---|---|---|---|
| Soft core minimization | 10000 | - | 10000 | 10000 | 10000* | 0 |
| Dry minimization | 2000 | - | 1000 | 1000 | 1000* | 0 |
| Solvent minimization | 5000 | - | 4000 | 2000 | 1000 | 1000 |
| Equilibration 1 | 125000 | 1 | 4000 | 2000 | 1000 | 1000 |
| Equilibration 2 | 125000 | 1 | 2000 | 1000 | 400 | 400 |
| Equilibration 3 | 125000 | 1 | 1000 | 500 | 400 | 200 |
| Equilibration 4 | 250000 | 2 | 500 | 200 | 200 | 200 |
| Equilibration 5 | 250000 | 2 | 200 | 50 | 40 | 100 |
| Equilibration 6 | 250000 | 2 | 50 | 0 | 0 | 0 |

* Position restraint for heavy atoms of POPC in central cavity

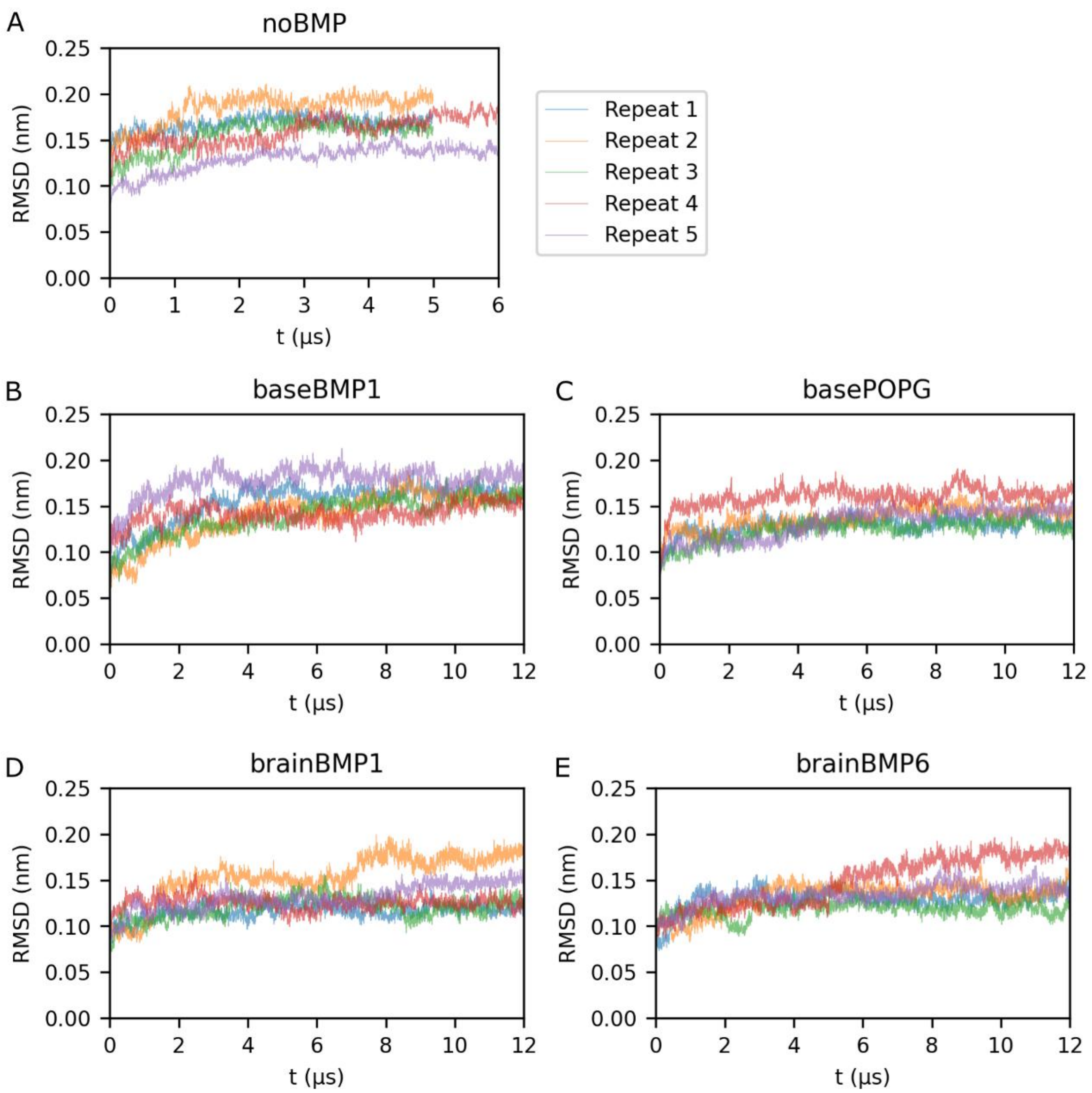


**Figure S1.** RMSD of the $c_8$-ring backbone of all trajectories from each simulation system: (A) noBMP, (B) baseBMP1, (C) basePOPG, (D) brainBMP1, (E) brainBMP6 with the trajectories from each repeat coloured as indicated by the legend.

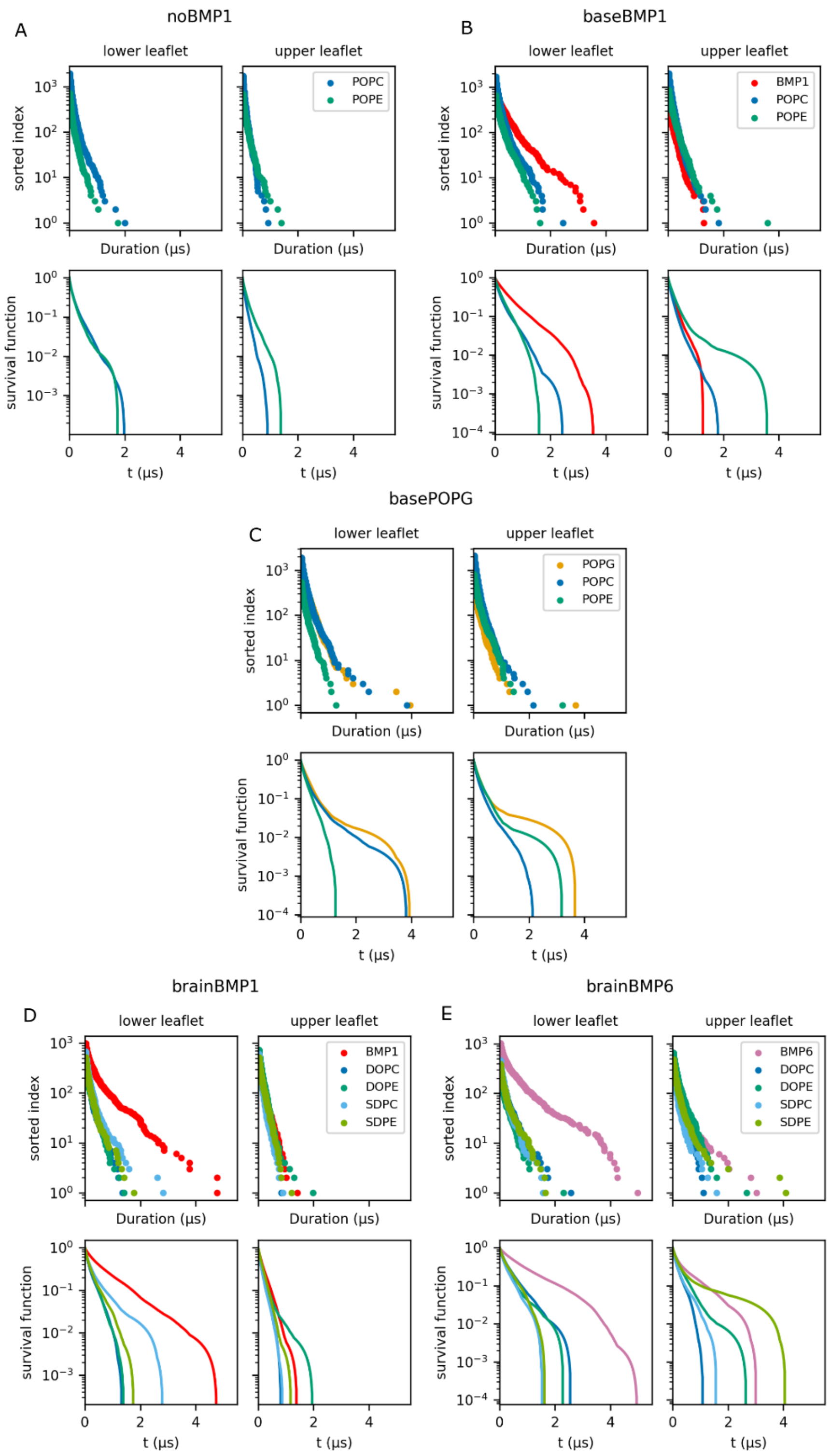


**Figure S2.** Log plots of contact durations and survival functions for all simulation systems for each lipid species per membrane leaflet. (A) baseBMP, (B) noBMP, (C) basePOPG, (D) brainBMP6, and (E) brainBMP1 membrane systems.

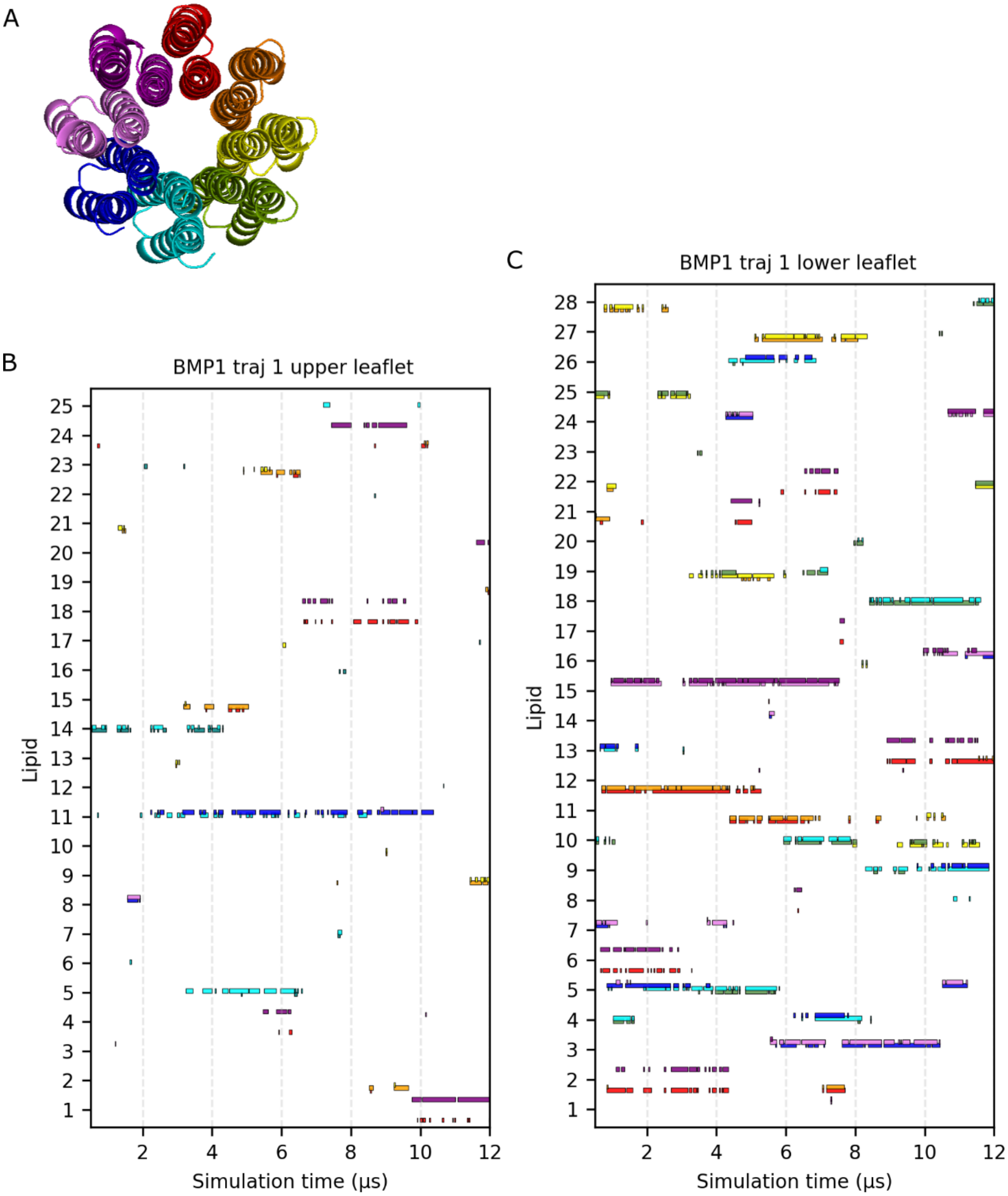


**Figure S3.** BMP1 lipid contacts with protein chains, for an example trajectory of the baseBMP system. (A) $c_8$-ring shown in cartoon representation coloured by chain. The same colour coding is used to indicate which chains a lipid is in contact with at time $t$ during the simulation. (B–C) Lipid contacts for BMP1 for the upper and lower membrane leaflets from a single trajectory.

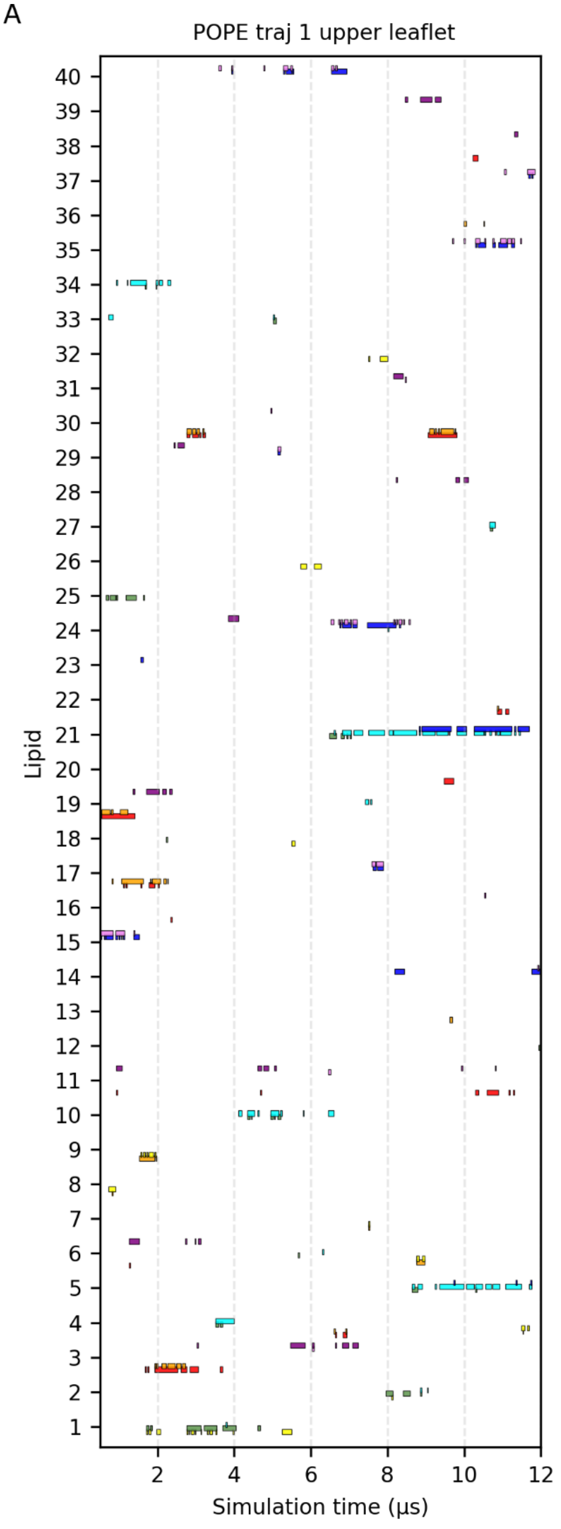


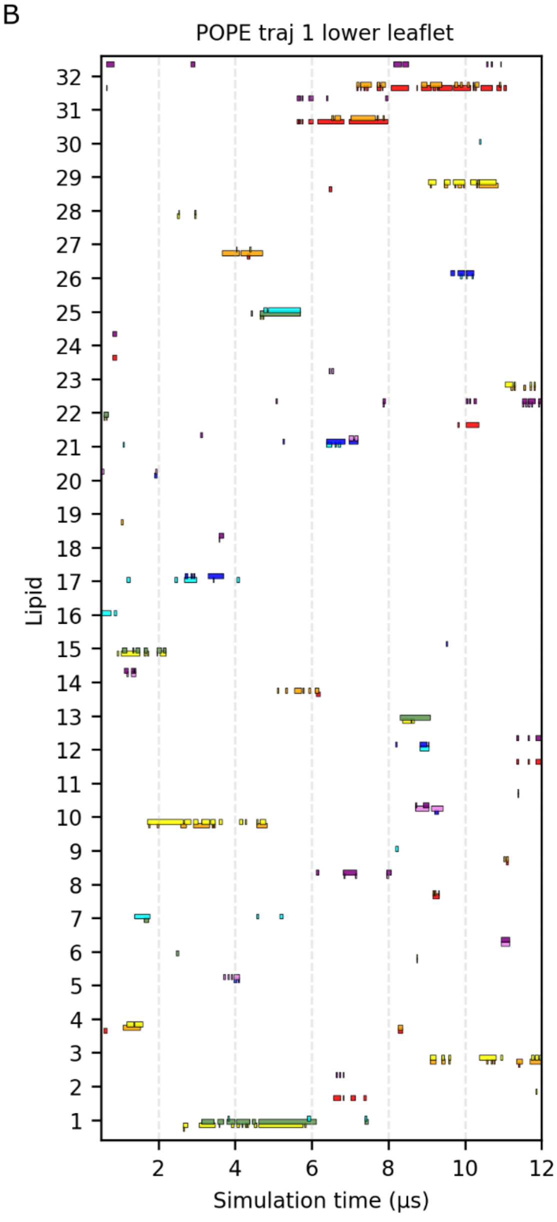


**Figure S4** POPE lipid contacts with protein chains for an example trajectory of the baseBMP system. Colouring follows the same scheme as in Figure S3. (A) for the upper leaflet, (B) for the lower leaflet.

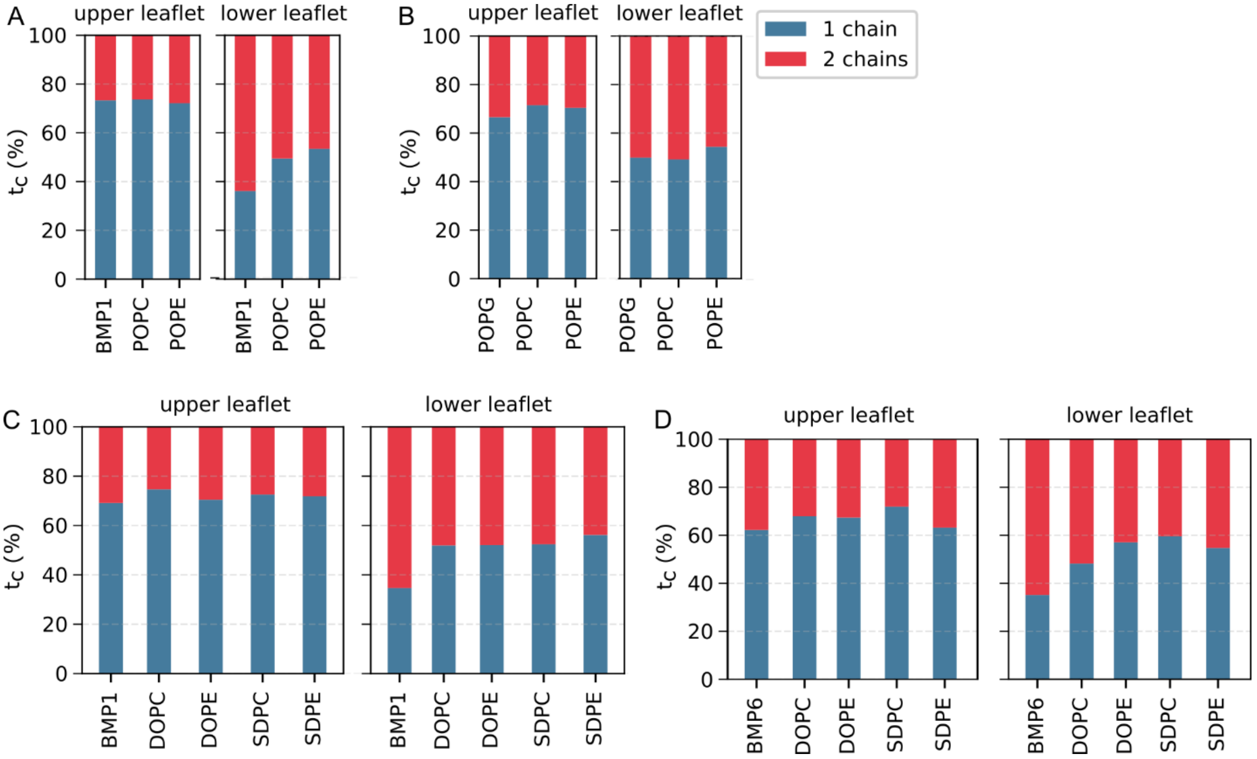


**Figure S5.** Contact time percentage $t_c$ per number of chains for each lipid species and membrane leaflet from the following membrane systems: (A) baseBMP, (B) basePOPG, (C) brainBMP6, and (D) brainBMP1.

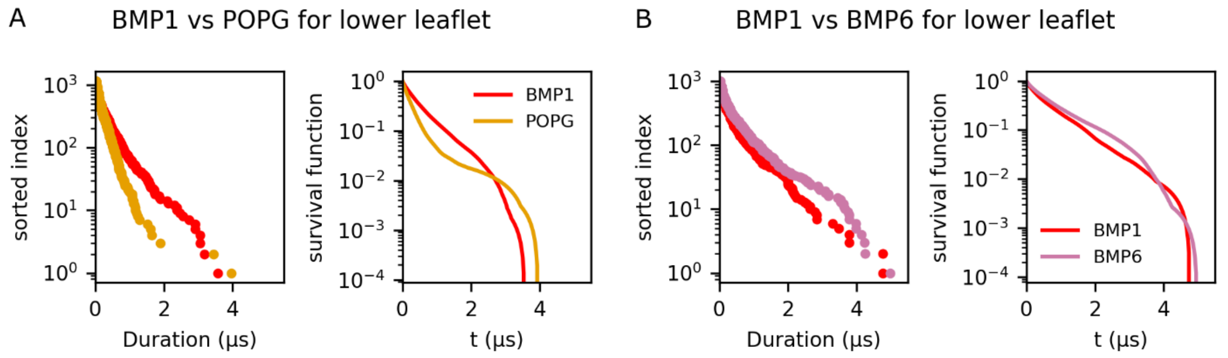


**Figure S6.** Comparison of contact-duration and survival-function distributions between anionic lipids in the lower membrane leaflet. (A) Comparison between POPG and BMP1 in ternary membranes containing POPC, POPE, and BMP1 or POPG. (B) Comparison between BMP1 and BMP6 in the brain-mimicking membrane systems (brainBMP1/brainBMP6).

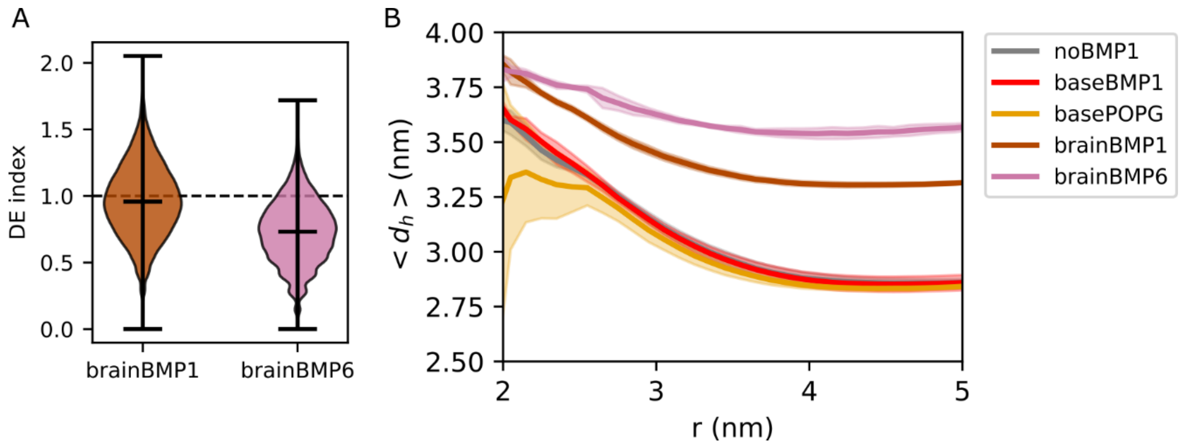


**Figure S7.** (A) Cholesterol DE index distributions for the BMP1- and BMP6-containing membrane systems shown as violin plots, with median, maximum, and minimum values shown in black. The dotted line indicates DE index = 1. (B) Membrane thickness as a function of distance from the protein for all simulated systems. Shaded areas indicate standard deviations obtained by averaging thickness profiles over individual simulation runs.
